# MAPK13 controls structural remodeling and disease after epithelial injury

**DOI:** 10.1101/2024.05.31.596863

**Authors:** Kangyun Wu, Yong Zhang, Dailing Mao, Courtney A. Iberg, Huiqing Yin-Declue, Kelly Sun, Shamus P. Keeler, Hallie A. Wikfors, Deanna Young, Jennifer Yantis, Stephen R. Austin, Derek E Byers, Steven L. Brody, Erika C. Crouch, Arthur G. Romero, Michael J. Holtzman

## Abstract

All living organisms are charged with repair after injury particularly at epithelial barrier sites, but in some cases this response leads instead to structural remodeling and long-term disease. Identifying the molecular and cellular control of this divergence is key to disease modification. In that regard, stress kinase control of epithelial stem cells is a rational entry point for study. Here we examine the potential for mitogen-activated protein kinase 13 (MAPK13) regulation of epithelial stem cells using models of respiratory viral injury and post-viral lung disease that resembles asthma. We show that *Mapk13* gene-knockout mice handle acute infectious illness as expected but are protected against basal-epithelial stem cell (basal-ESC) hyperplasia and mucous metaplasia. In corresponding organoid models, *Mapk13*-deficiency directly controls the basal-ESC program for stemness endpoints of hyperplasia and mucous metaplasia. Extension to human studies shows induction/activation of MAPK13 in basal-epithelial cells in lung tissue samples from asthma and COPD patients. Further, in human organoid models, *MAPK13* mRNA knockdown regulates basal-ESC stemness similarly to mouse models. Together, the data identify MAPK13 as a control point for structural remodeling after epithelial injury and a suitable target for down-regulation as a disease-modifying strategy.

**New and noteworthy:** This study identifies a distinct role for stress kinase MAPK13 in controlling the epithelial stem cell response to injury and the consequent development of structural remodeling. The present model has direct implications for lung injury and subsequent disease triggered by respiratory viruses and other inhaled hazards. The findings also refine a hypothesis for therapeutic intervention based on proper scaling of MAPK13 function including down-regulation with selective kinase inhibitors.

## Introduction

Organisms are constantly challenged with a wide spectrum of injuries and the need for a repair process to return to proper physiological function. While this goal is often achieved, in some cases, the process results in a skewed response that ends in structural remodeling and dysfunction of the injured tissue site. Therefore, it is key to define the cellular and molecular controls of these outcomes to foster and preserve proper repair versus remodeling and abnormal function. This paradigm is particularly common at epithelial barrier sites wherein epithelial stem cells and stress-related signaling molecules orchestrate the response to injury from a host of environment agents. Notable among these molecules are an overlapping and interacting set of stress kinases presumably designed for redundancy and diversity. Thus, the complexity of this network has made it difficult to assign function to specific kinases and ultimately to modify kinase behavior to preserve or enhance repair and prevent or reverse abnormal remodeling and disease. Thus, a first step remains to better identify stress kinase control of the epithelial repair and remodeling process after injury from inhaled agents.

Among the agents for study, the respiratory barrier is especially exposed to inhaled toxins and infectious agents with every breath and therefore maintains an active sentinel system to restore integrity at the barrier site. Here again, however, the normal program for recovery can be skewed towards structural remodeling and chronic respiratory disease. This divergence is well established after respiratory viral infection can progress to post-viral lung disease manifest as airway inflammation and mucus production that is characteristic of asthma and an overlapping subset of chronic obstructive pulmonary disease (COPD) (1–11). In that context, basal-epithelial stem cells (basal-ESCs) represent a stereotyped feature of the epithelial barrier program. This stem cell population can be reprogrammed for hyperplasia and metaplasia that disrupts lung function in experimental mouse models of post-viral asthma (3, 12–14). A similar basal-epithelial cell response is also found after severe respiratory viral infection in humans (4, 15, 16) but is not yet linked functionally to post-viral lung disease of any kind, including the development of post-viral asthma (17).

Relatedly, the molecular control over basal-ESCs that determines post-viral repair versus remodeling is also uncertain. One of the most studied molecular responses to injury are stress kinases, particularly the mitogen activated protein kinase (MAPK) family. The original family member MAPK14 (aka MAPK-p38α) was proposed as a master regulator of the inflammatory response to tissue damage (18). Unfortunately, this candidate selection has not resulted in modifying the response to injury in the lung or other tissue sites (19–21). As an alternative, our studies of basal-ESCs cultured from human airways identified MAPK13 (aka MAPK-p38δ) as a dominant control point for basal-ESC differentiation to mucous cells (1). This finding was intriguing given the primordial role of mucus production in host defense and pathogenic role of excess mucus production in virus-triggered diseases such as asthma and COPD (1, 3, 7, 22, 23). The fundamental nature of MAPK13 control also derives from studies of diabetes, sepsis, and cancer (24–29). Given this diversity, we questioned here whether MAPK13 might represent a fundamental checkpoint for epithelial stem cell reprogramming towards disease endpoints. The present study therefore addresses whether MAPK13 might be essential for basal-ESC reprogramming using well-controlled gene knockout or mRNA knockdown approaches in models of the epithelial responses to viral infection in vivo in mice and ex vivo in mouse and human airway organoids (1–3, 14, 30–40). Together, the experimental approaches provide a paradigm for epithelial barrier cells in balancing tissue repair versus structural remodeling and thereby delivering MAPK13 as a precise target for a disease-modifying strategy.

## Results

### MAPK13 controls post-viral lung disease

As introduced above, MAPK13 function was implicated in mucous metaplasia in human airway epithelial cells in air-liquid interface culture (1). However, any corresponding role for MAPK13 alone in regulating mucus production in vivo or other aspects of basal-epithelial stem cell (basal-ESC) reprogramming (particularly cell stemness) in any model still needed to be better defined. To address these issues, we engaged a mouse model that resembles lung diseases (asthma and overlap with COPD) that are linked to respiratory viral infection in humans (1–4). In this model, infection with a natural parainfluenza-virus pathogen known as Sendai virus (SeV) manifests a typical proximal-to-distal pattern of respiratory infection followed by a remarkably long asthma endotype/phenotype validated in humans (1–3, 14, 30–43). To assess MAPK13 function in this model, we generated *Mapk13^−/−^* mice (**Supplemental Figure 1A,B**) that manifested slightly attenuated acute infectious illness to wild-type (WT) mice as tracked by less weight loss and similar lung levels of viral RNA (**Figure 1A,B**). Despite clearance of infectious virus, lung levels of MAPK13 continued to increase during the development of long-term lung disease based on immunoblotting (**Supplemental Figure 1C**). This increase coincided with increased immunostaining for MAPK13^+^KRT5^+^ basal-epithelial cells that was attenuated in *Mapk13*^−/−^ mice **(Figure 1C,D).** MAPK13 was primarily nuclear in KRT5^+^ basal-epithelial cells, as expected for kinase phosphorylation/activation. These changes were associated with attenuation of periodic acid Schiff (PAS) and hematoxylin tissue staining (reflecting mucus production and cellularity, respectively) in lung sections at 49 d after SeV infection when tissue remodeling and disease phenotypes are otherwise maximal (**Figure 1E**). The attenuation of basal-epithelial cell hyperplasia/metaplasia and histopathology in *Mapk13*^−/−^ mice compared to WT control mice was significant based on quantitative morphology (**Figure 1F**). Relatedly, we found significant down-regulation of the usual increase in AQP3^+^ basal-epithelial cells (but not AQP3^−^ non-basal epithelial cells) in *Mapk13*^−/−^ mice (**Figure 1G,H**). This finding was consistent with attenuation of basal-epithelial cell but not alveolar-epithelial type 2 (AT2) cell hyperplasia in *Mapk13*^−/−^ mice.

**Fig. 1.**
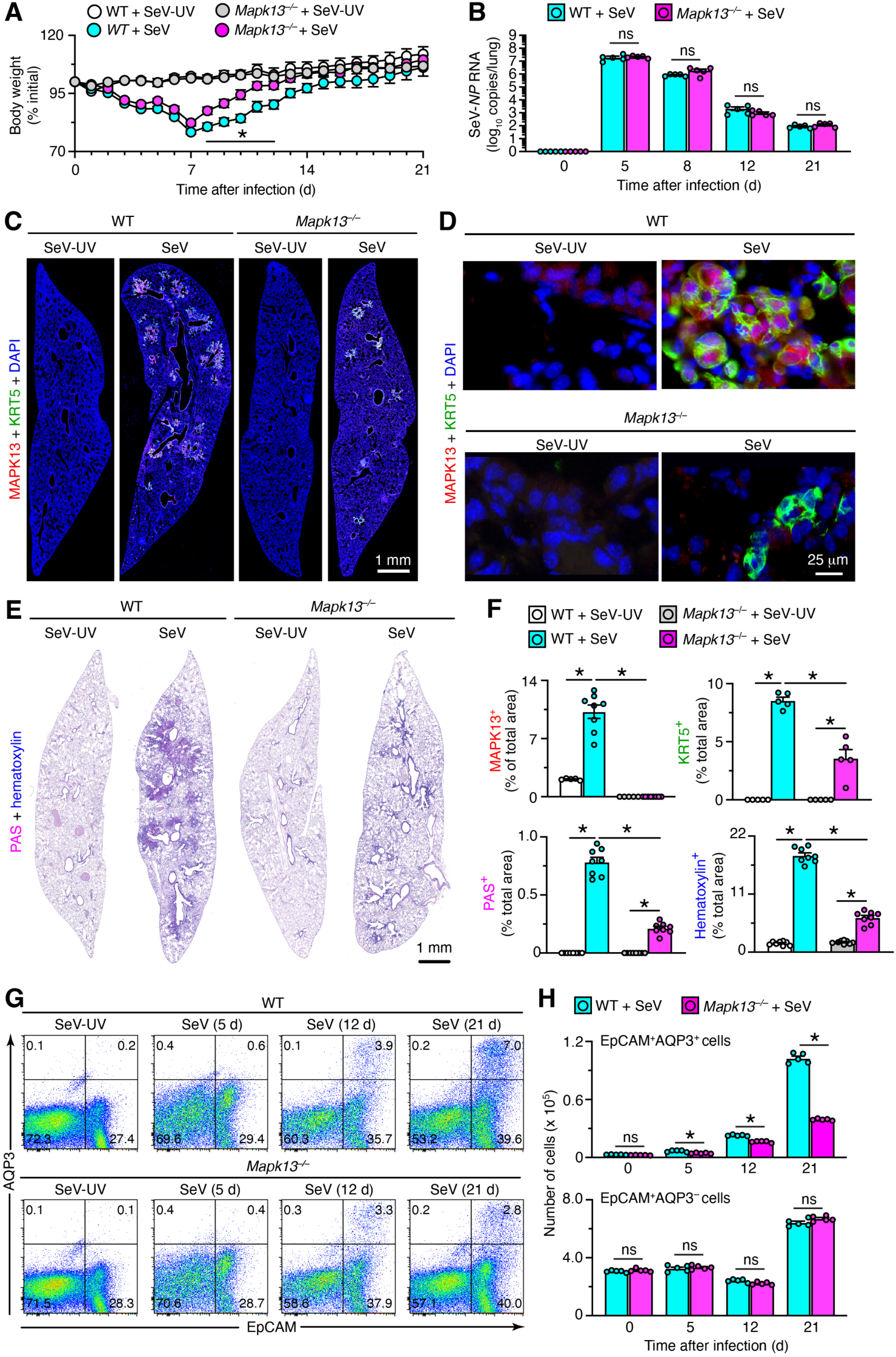
*Mapk13* gene-knockout attenuates basal-ESC hyperplasia/metaplasia in a mouse model of post-viral asthma using SeV infection. **A**, Body weights for wild-type (WT) and *Mapk13*^−/−^ mice after infection with Sendai virus (SeV) or SeV-UV control. **B,** SeV-*NP* RNA levels in lung tissue from WT and *Mapk13*^−/−^ mice after SeV or SeV-UV. **C**, Immunostaining for MAPK13 and KRT5 with DAPI counterstaining in lung sections from WT and *Mapk13*^−/−^ mice at 49 d after SeV or SeV-UV. **D**, Higher magnification of basal-epithelial cell remodeling regions for conditions in (C). **E**, PAS and hematoxylin staining for conditions in (C). **F**, Quantitation of staining for (C.D,E). **G**, Flow cytograms of lung epithelial cells (CD31^−^CD45^−^EpCAM^+^Aqp3^+^ and CD31^−^CD45^−^EpCAM^+^Aqp3^−^) at 5, 12, and 21 d after SeV or SeV-UV. Values indicate the percentage of cells within each gate. **H**, Cell numbers for conditions in (G). Values represent mean ± s.e.m. (n=5-10 mice per condition). \**P* <0.05 by ANOVA and Tukey correction; ns, not statistically significant.

In concert with attenuation of remodeling and histopathology, *Mapk13*^−/−^ mice also showed significant down-regulation of specific biomarkers designed to track disease endpoints based on lung tissue mRNA levels. This approach showed that *Mapk13*^−/−^ mice exhibited significant attenuation of biomarkers for basal-ESC hyperplasia marked by *Krt5, Aqp3*, and *Trp63*; alarmin signal marked by *Il33*, immune activation marked by *Serpinb2*, *Ltf, Cxcl17,* and *Nos2*; type 2 inflammation marked by *Il13, Arg1, Trem2,* and *Spp1,* and type 1/2 inflammation marked by *Il6* (**Figure 2A-E**). We found no significant induction or blockade of conventional type 1 inflammation marked by *Ifng, Tnfa,* and *Il1b* mRNA levels (**Figure 2F**), consistent with data that this signal (that is often MAPK14-dependent) is not a primary driver for chronic disease after viral infection in the present model (1, 44, 45). As expected, *Mapk13* gene induction/expression was selective (relative to related *Mapk11*, *12*, and *14*) and was not detected in *Mapk13*^−/−^ mice (**Figure 2G**). The most effective attenuation was found for mucous metaplasia marked by *Muc5ac* and *Clca1*(**Figure 2H)**. The *Muc5b* mRNA signal for mucous metaplasia was relatively low (**Figure 2H**), but this finding might be due to probe sensitivity or post-transcriptional regulation given higher signal found by immunostaining. Mucous metaplasia developed in concert with reciprocally lower levels of club cells marked by *Scgb1a1* mRNA (**Figure 2I**).

**Fig. 2.**
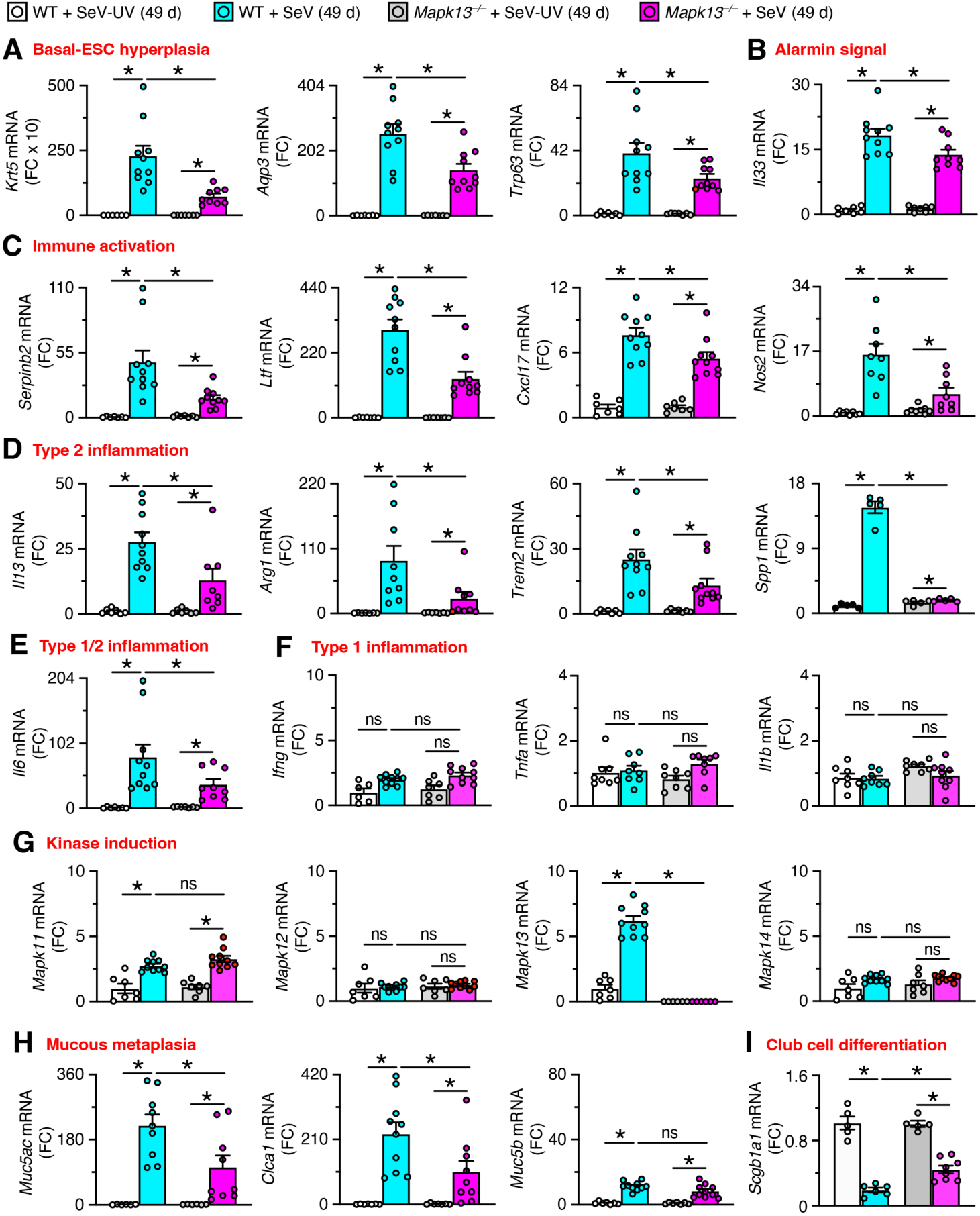
*Mapk13* gene-knockout attenuates mRNA biomarkers in a model of post-viral asthma. **A-I**, Lung tissue levels of mRNA biomarkers in WT and *Mapk13*^−/−^ mice at 49 d after SeV or SeV-UV to track basal-ESC hyperplasia (A), alarmin signal (B), immune activation (C), type 2 inflammation (D), type 1/2 inflammation (E), type 1 inflammation (F), kinase induction (G), mucous metaplasia (H), and club cell differentiation (I). Values represent mean ± s.e.m. (n=5-10 mice per condition). \**P* <0.05 by ANOVA and Tukey correction.

In line with the effects on mRNA biomarkers in *Mapk13*^−/−^ mice, we also found correction of clinical disease phenotypes. These included attenuation of immune activation based on NOS2^+^ immunostaining (**Figure 3A**), macrophage level based on F4/80^+^ immunostaining (**Figure 3B**), eosinophil infiltration based on major basic protein (MBP) immunostaining (**Figure 3C**), and mucous metaplasia based on immunostaining for MUC5AC and MUC5B (**Figure 3D**). Each of these findings was significant based on quantitative morphology (**Figure 3E-H**). To further define whether the downregulation of these disease biomarkers translated to improvements in mucus production and pulmonary function, we also checked levels of mucus production in bronchoalveolar lavage fluid along with oxygenation and airway reactivity that are also regularly associated with airway disease. Notably, we found attenuation of mucus production marked by MUC5AC and CLCA1 levels (**Figure 3I**), hypoxemia monitored by SpO_2_ using oximetry (**Figure 3J**), and airway hyperreactivity assessed using inhaled methacholine challenge (**Figure 3K**). Together, these findings support the proposal that MAPK13 deficiency alone can attenuate excessive basal-ESC hyperplasia, immune activation, and mucous metaplasia to improve pulmonary function.

**Fig. 3.**
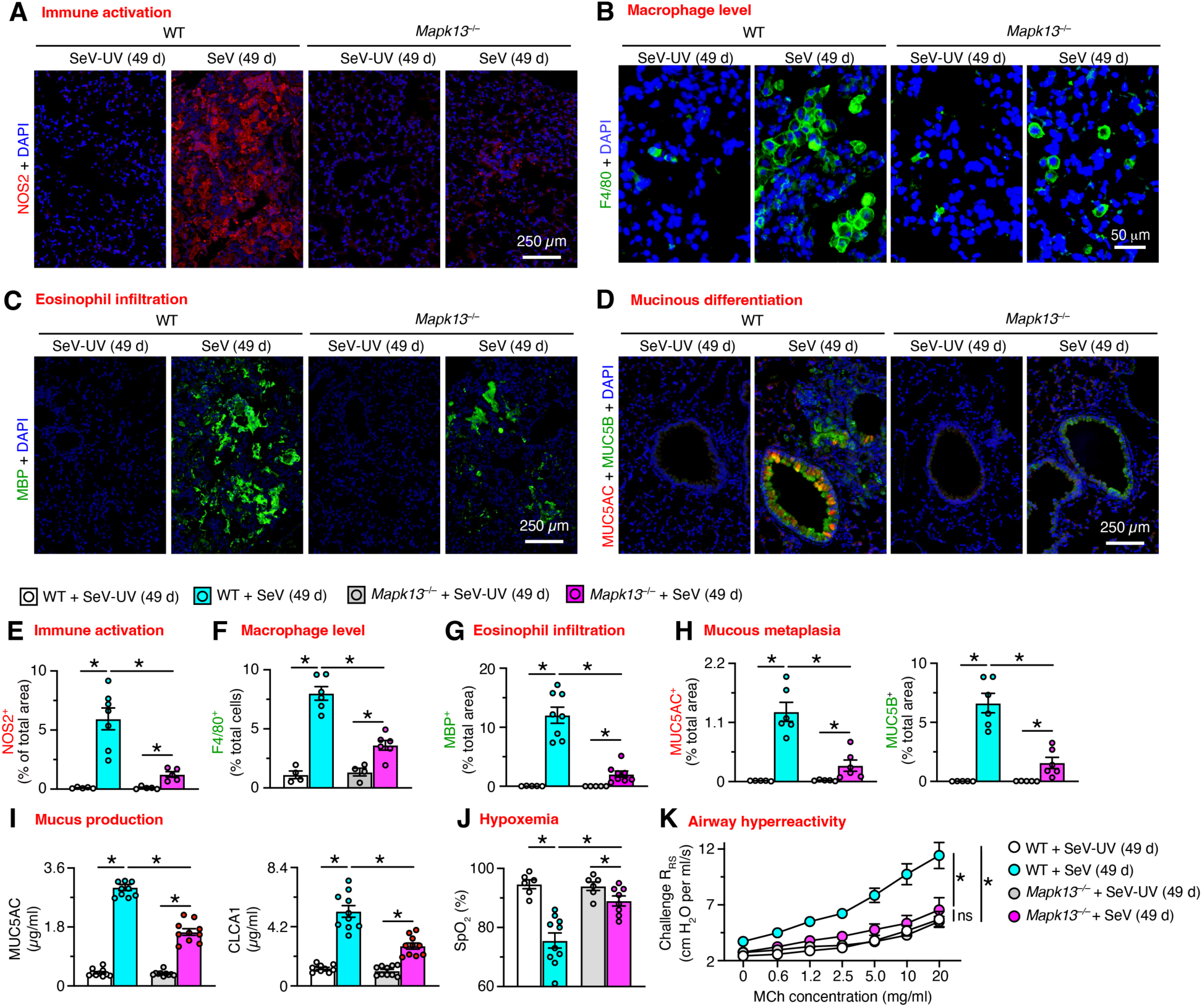
*Mapk13* gene-knockout attenuates clinical readouts in a model of post-viral asthma. **A**, Immunostaining for NOS2 with DAPI counterstaining of lung sections from WT and *Mapk13^−/−^* mice at 49 d after SeV or SeV-UV. **B**, Immunostaining for F4/80 with DAPI counterstaining for conditions (A). **C**, Immunostaining for major basic protein (MBP) with DAPI counterstaining for conditions (A). **D,** Immunostaining for MUC5AC and MUC5B with DAPI counterstaining for conditions in (A). **E-H**, Quantitation of immunostaining for (A-D). **I**, Levels of MUC5AC in bronchoalveolar lavage fluid for conditions in (A). **J**, Oximeter levels for blood oxygen saturation (SpO_2_) for conditions in (A). **K,** Values for respiratory system resistance (R_RS_) at baseline and after inhaled methacholine (MCh) for conditions in (A). Values represent mean ± s.e.m. (n=5-9 mice per condition). \**P* <0.05 by ANOVA and Tukey correction.

### MAPK13 directly controls basal-ESC function

We next aimed to extend our experimental findings in mouse models to humans and better establish a direct role for MAPK13 in control of basal-ESC function. To start, we assessed whether MAPK13 expression and activation might be linked to basal-ESC reprogramming in human lung disease. Given the challenge of obtaining lung tissue samples in the setting of severe viral infection, we characterized lung tissue samples from asthma and COPD patients in our registry (see **Table 1** for clinical characteristics). Previously, we found that these samples exhibit a post-viral phenotype that includes overexpression of alarmin (IL-33), type 2 inflammation (IL-13), and mucus production (1, 3). Here we found increases in IL-33 and MAPK13 expression localized to KRT5^+^ basal-epithelial cells particularly in asthma and COPD patients (**Figure 4A**). MAPK13 expression was often further localized to a nuclear site, consistent with phosphorylation/activation and nuclear translocation that is typical of MAPK and kinase signaling in general (45).

**Fig. 4.**
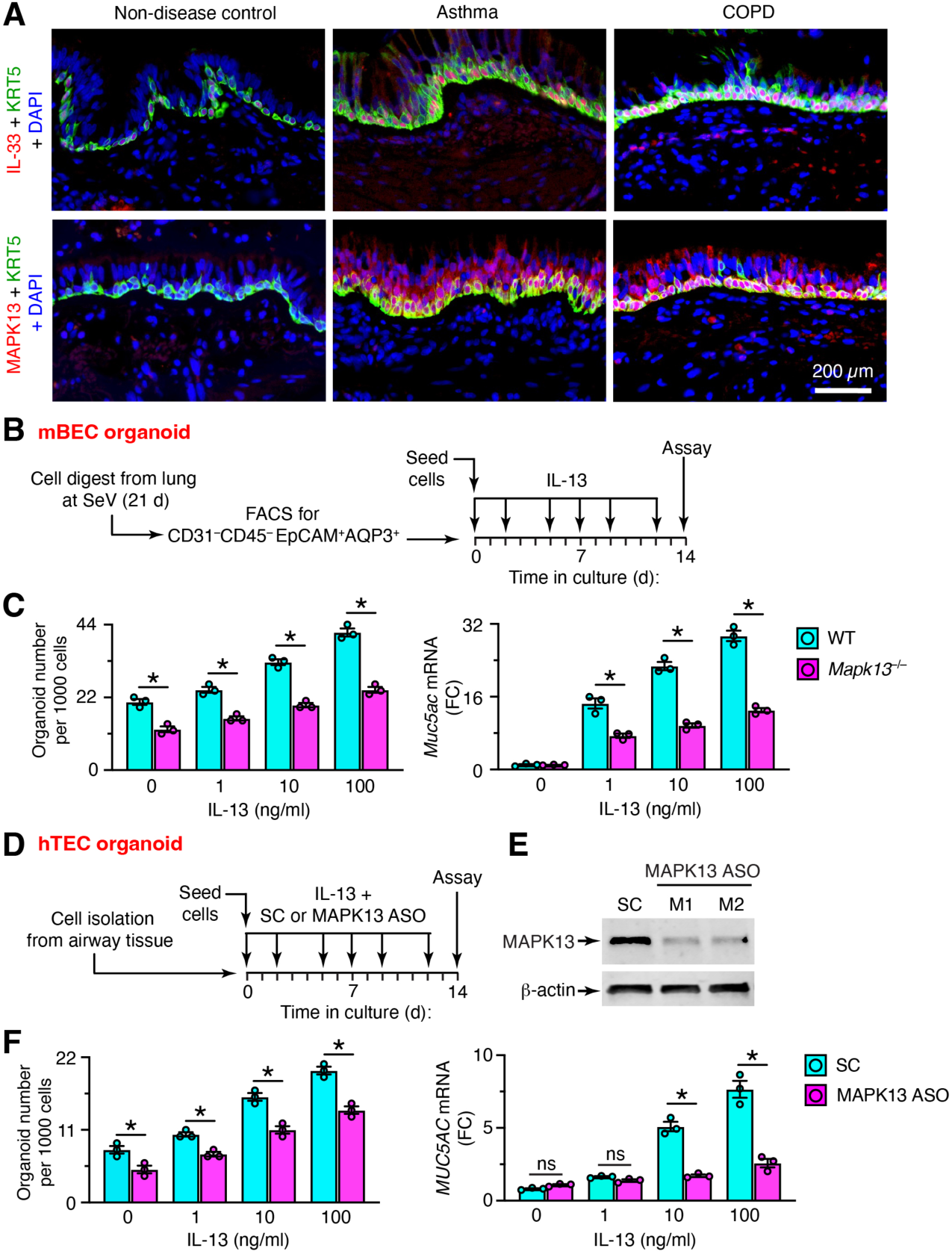
MAPK13 is activated in human airway disease and is directly required for basal-ESC stemness and mucous metaplasia. **A**, Representative immunostaining for IL-33 plus KRT5 or MAPK13 plus KRT5 with DAPI counterstaining in lung sections from non-disease control (n=11), asthma (n=10), and COPD (n=10) subjects (inset, 5x). **B**, Protocol for mouse bronchial-epithelial cell (mBEC) derived organoids established with CD45^−^CD31^−^ EpCAM^+^AQP3^+^ cells isolated from lungs of WT and *Mapk13*^−/−^ mice at 21 d after SeV infection and then cultured with and without IL-13 at indicated concentrations. **C**, Levels of cell stemness (marked with organoid formation) and mucous metaplasia (marked with *MUC5AC* mRNA) for conditions in (B). Values represent mean ± s.e.m. (n=5-9 mice per condition). **D,** Protocol for human tracheobronchial epithelial cell (hTEC) derived organoids with *MAPK13* mRNA knockdown using culture conditions for (B) plus treatment with scrambled control (SC) or anti-sense oligonucleotide (ASO). **E,** Western blot for hTEC treatment with SC or ASOs (designated M1 and M2) using conditions in (D). **F**, Levels of cell stemness for conditions in (D) with SC and M1 ASO treatment. Values represent mean ± s.e.m. for a single subject representative of 3 subjects. \**P* <0.05 by ANOVA and Tukey correction.

To next address MAPK13 control of cell function, we extended our analysis of mouse and human samples to airway organoids derived from basal-ESCs as a new approach methodology. For mice studies, we established conditions for generating airway organoids from WT and *Mapk13^−/−^* mice and maintaining the model with and without IL-13 to mimic in vivo conditions (**Figure 4B**). This system showed that baseline and IL-13-stimulated organoid formation as well as IL-13-stimulated mucous metaplasia were downregulated with *Mapk13* deficiency (**Figure 4C**). For human studies, we established airway organoids with primary human tracheobronchial epithelial cells (hTECs) and achieved MAPK13 knockdown using a newly applied chemical method for anti-sense oligonucleotide (ASO) to target *MAPK13* mRNA (**Figure 4D**) and thereby decrease MAPK13 protein (**Figure 4E**). Similar to the pattern in the mouse organoid model, we found significant attenuation of basal-ESC stemness (tracked with organoid formation) and even more effective blockade of mucous metaplasia under these conditions (**Figure 4F**). Together, these results provided evidence that MAPK13 is overexpressed, activated, and susceptible to targeted blockade of disease-related function in basal-ESCs as predicted from our viral-infection mouse model.

## Discussion

In this study, we identify MAPK13 as a critical check point for structural remodeling and chronic lung disease after respiratory viral infection. Key findings include: (1) *Mapk13* gene knockout alone is sufficient to attenuate basal-ESC reprogramming for hyperplasia/metaplasia, immune activation, and mucous metaplasia in a mouse model of epithelial injury and post-viral lung disease; and (2) *MAPK13* mRNA knockdown similarly downregulates basal-ESC stemness and mucous metaplasia in corresponding mouse and human airway organoid models. Mouse studies localize MAPK13 induction and activation to basal-epithelial cells in situ at sites of disease, and human studies detect similar induction/activation in asthma and COPD. The data thereby define MAPK13 as a practical target for modifying the structural remodeling and dysfunction that can develop after epithelial injury due to severe respiratory viral infection as diagrammed in **Figure 5**. Here we highlight three aspects of this research progress.

**Fig. 5.**
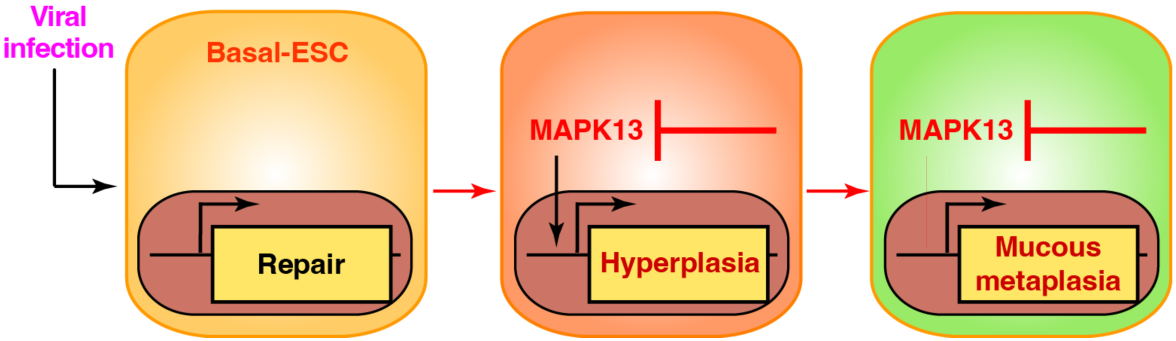
Scheme for MAPK13 control of basal-ESC reprogramming as found in post-viral asthma. Respiratory viral infection can trigger basal-ESC activation for regeneration and repair to restore homeostatic conditions. However, basal-ESCs can also be reprogrammed to progress to hyperplasia and mucous metaplasia. Direct MAPK13 control of these basal-ESC endpoints is established in the present data using mouse and human models.

First, an important premise of the present study was that the definition of MAPK13 function remained incomplete despite previous studies of MAPK13 deficiency in gene-knockout mice. Some of the uncertainty was due to the frequent performance of MAPK13 studies in combination with MAPK12 deficiency, based on the similar protein sequence/structure and in some cases tissue distribution of these two potentially redundant kinases (28, 46–52). Given the present results, it is rational to consider the function of MAPK13 on its own. In that regard, MAPK13 solo function has been detected in mouse models of neutrophil infiltration in LPS-induced lung injury (26), pancreatic beta-cell secretion of insulin (25), LPS-induced lung injury (26), and keratinocyte hyperplasia after carcinogenic stimulation (27). However, these previous studies did not precisely localize MAPK13 expression and activation as done in the present analysis using tissue immunostaining and corresponding cell isolation. Thus, the site of action for MAPK13 was generally inferred from related ex vivo experiments. Despite this drawback, the overlap of the present findings with basal-epithelial cell hyperplasia after epithelial injury in the lung appear to overlap with a similar phenotype in skin and the related theme of epithelial stemness. This connection further reinforces the proposal for a role for MAPK13 in the jump from epithelial barrier repair to structural remodeling after injury. It is also possible that MAPK13 might coordinate basal-ESC hyperplasia with immune activation and mucous metaplasia based on downstream signal transduction to kinase and chromatin targets, but these connections still need to be defined.

Second, and relevant to the spectrum of stress kinase function, the present findings come in the context of our previous work on MAPK13 in human cell and mouse models of chronic lung disease and corresponding analysis of lung tissue samples from this type of disease. These earlier studies first uncovered a role for MAPK13 in controlling basal-ESC differentiation to mucous cells in concert with MAPK13 induction/activation in chronic respiratory disease (1). In this initial work, definition of function relied on study of human basal-ESC cultures subject to siRNA-mediated gene-knockdown or treatment with a relatively weak MAPK13 inhibitor (NuP-43; BIRB-796) and first-generation compounds that improved MAPK13 enzyme inhibition but maintained broad kinase-blocking activities (1). Subsequently, this approach was extended to a second-generation MAPK13-14 inhibitor (designated NuP-3) that was also effective in blocking mucus production in hTEC cell cultures and that also prevented airway inflammation and mucus production in minipig models of disease using cytokine-challenge and viral-infection (44). However, these models manifested only short-term inflammation and mucus production without the full features of long-term basal-ESC hyperplasia/metaplasia found in the presently used mouse model and in humans (3, 4, 14, 35, 36). The present work advances these previous approaches by showing that MAPK13 alone can also control basal-ESC function based on more specific techniques for MAPK13-deficiency in vivo in mice and in vitro in mouse and human basal-ESC-derived organoids. The findings thereby suggest a more fundamental role for MAPK13 in the response to epithelial barrier injury and an opportunity to improve barrier regeneration guided by the present lung function readouts for mucus production, blood/tissue oxygenation, and airway reactivity.

With this advance, we expect the present data to provide an alternative to the canonical focus on conventional stress kinases in inflammatory disease, including lung disease. More specifically, there has been primary attention to the related kinase MAPK14 as a functional and therapeutic target of interest (19, 53–63). Indeed, in models of inflammatory disease, MAPK14-specific inhibitors are highly effective in blocking cytokine (e.g., TNF-α and IL-1β) production and/or signal transduction (64). This mechanism predicted therapeutic benefit in a variety of conditions including acute and chronic respiratory diseases (65). However, even the most advanced versions of these candidate drugs have not proved effective in clinical trials of patients with chronic lung disease due to COPD where these types of cytokines are found at higher levels (57). The data here suggest a basis for this failure by implicating the independent action of MAPK13 in a disease process that is instead linked to type 2 inflammation. This connection is illustrated by the present biomarker panel that features this alternative type of inflammation (marked especially by cytokine IL-13 expression). In contrast, the biomarkers for conventional inflammation (TNF-α and IL-1β) showed no significant induction in the present model of structural remodeling and chronic lung disease that developed after viral infection.

Third, despite the MAPK13 link to epithelial cells, the present data does not exclude an additional role for MAPK13 function in immune cells, particularly those participating in the type 2 immune response as a stereotyped feature of tissue repair and remodeling. In that regard, we provided evidence of monocyte-derived dendritic cell (moDC) action upstream along with macrophage, innate lymphoid cell type 2 (ILC2), and NKT cell action downstream of basal-ESC activation (2, 34, 37, 38). These epithelial-immune cell collaborations thereby provide a feed-forward mechanism for the observed combination of disease phenotypes (basal-ESC hyperplasia/metaplasia, type 2 inflammation, and excess mucus production). These phenotypes are characteristic of post-viral lung disease in mice as a model of asthma and overlapping airway disease in COPD in humans guided by the present biomarker panel. Defining the precise role of each epithelial-stem cell and immune-cell population is a goal of future studies. However, conditional knockout strategies for basal-ESCs can change baseline conditions (14, 66) when targeting MAPK13 and related members of the CMGC kinase superfamily (29, 67). Further, selective down-regulation of specific myeloid cell subsets is still imprecise with existing tools. Further, few if any of these genetic approaches can be selective for the lung, which results in off-target effects that complicate practical application and interpretation. This matrix underlines the need for small-molecule kinase inhibitors that can selectively engage the target molecule (MAPK13) and tissue (airway) as provided in our initial report (68). These strategies can be complemented using new approach methodologies to human cell/organoid culture asperformed in the current report.

In sum, the present data solidify the proposal for a primordial response to epithelial injury that depends on MAPK13 and suggest a MAPK13-targeted strategy to improve this response if it skews to structural remodeling and disease. The results also highlight one of the most common types of epithelial injury in the form of respiratory viral infection. Significant evidence demonstrates that this same pattern of epithelial and immune cell responses develops after injury due to a broad spectrum of respiratory viruses (including human enterovirus, influenza virus, respiratory syncytial virus, and human rhinovirus) and other inhaled agents (such as airborne pollutants) (35, 36, 69–71). In each case, there is experimental and clinical evidence for initiation, progression, and/or exacerbation of chronic respiratory disease. The tissue distribution of MAPK13 suggests that it might provide a similar function at other sites and in other forms of tissue injury. Future studies should provide a coherent picture of epithelial injury, remodeling, and corresponding therapy using the present findings as a more specific guideline for a correctable target kinase.

## Materials and Methods

### Mouse models

Male and female wild-type (WT) C57BL/6J mice (000664) mice were obtained from Jackson Laboratory. The *Mapk13* gene knockout mice (*Mapk13^−/−^*) were generated in the C57BL/6J background using CRISPR/Cas9 technology with gRNAs that were designed to target early in exon 1 based on a sequence to minimize any off-target effect. Synthetic gRNAs were purchased from IDT Technologies and complexed with recombinant Cas9 protein before nucleofection in Neuro 2a cells for validation. The selected gRNA (tacctgccgcccgcgcacg) cleaved 8 nucleotides (cgtcggca) that resulted in a reading frame shift and stop codon introduction in exon 2 of the *Mapk13* gene. The ribonucleoprotein complex containing Cas9 protein (1 µg/µl) and gRNA (0.3 µg/µl) was then electroporated into single-cell embryos of C57BL/6J mice, and 25–30 eggs were transferred into each pseudo-pregnant female to generate founder mice. Knockout founders were identified by analyzing the PCR-amplified target region (forward primer 5′-ggaacgtacctgggcgag and reverse primer 5′-gtcccacgaactccgagatc) using next-generation sequencing to identify out-of-frame indels. Heterozygous F1 mice were obtained from matings between founders and WT mice, and homozygous F2 mice were obtained through sibling matings of F1 mice. The resulting *Mapk13*^−/−^ mice were found to reproduce and develop normally relative to WT control mice.

All mice were maintained and co-housed in a barrier facility using cages fitted with micro-isolator lids. Animal husbandry and experimental procedures were approved by the Animal Studies Committees of Washington University School of Medicine in accordance with the guidelines from the National Institutes of Health. Sendai virus (SeV, Sendai/52 Fushimi strain, ATCC VR-105) was obtained from ATCC and prepared and titered by plaque-forming assay and qPCR assay as described previously (2). Mice were infected with SeV (2.6 x 10^5^ PFU) as described previously (36). Virus or an equivalent amount of UV-inactivated virus or PBS alone was delivered intranasally in 30 µl of PBS under ketamine/xylazine anesthesia at 6-9 wk of age. Results from male and female mice were pooled since no significant differences were found between sexes as reported initially (72) and confirmed recently (36) and in the present experiments (data not shown). Viral titers for stock solutions and lung infections were monitored by quantitative PCR (qPCR) assay using primers for SeV-*NP* RNA as defined previously and in **Supplemental Table 2** using *SeV-NP*-expressing plasmids as an internal standard (36)

### Immunoblotting

For western blotting, lung tissue or cultured cells were lysed in RIPA buffer (ThermoFisher) supplemented with PhosSTOP (Sigma) and cOmplete PIC (Sigma) for 30 min at 4 °C. Tissue was homogenized in bead tubes and cell lysates were sheared 7 times with a 29G needle, and all samples were centrifuged at 14,100 x g at 4 °C. The supernatant was aliquoted and stored at -80°C until needed. Protein concentration was determined with the bicinchoninic acid (BCA) protein assay kit (ThermoFisher). Lysate (40-50 µg) was loaded into 4-15% Mini-PROTEAN® TGX gels (Bio-Rad) and separated by SDS-PAGE electrophoresis. Proteins were transferred to nitrocellulose membranes using a Trans-Blot® Turbo blot transfer pack (Bio-Rad). Nitrocellulose membranes were blocked with Odyssey blocking buffer (Li-COR Biosciences) and then primary antibodies for MAPK13 (AF1519, R&D), β-actin (ab8224, Abcam), or Gapdh (MAB374, Sigma) were incubated overnight at 4 °C in Odyssey blocking buffer with 0.2% Tween-20. Membranes were washed with 0.1% PBS-Tween-20 and then incubated with anti-mouse or anti-rabbit secondary antibodies (Li-COR) in Odyssey blocking buffer with 0.2% Tween-20 for 1 at 25 °C. Membranes were washed with 0.1% PBS-Tween-20, and protein bands were detected using an Odyssey CLx infrared imaging system (LI-COR Biosciences).

### Flow cytometry and FACS

Single cell suspensions were generated from minced lung tissue that was subjected to collagenase (Liberase TM Research Grade, Roche), hyaluronidase (Sigma), DNAse I (Sigma), and Dispase II (Roche) digestion for 45 min at 37 °C and then treated with ACK buffer (Lonza) to remove red blood cells. Following FcR blockade, lung cell suspensions were incubated with labeled antibodies and were sorted using a Sony SY3200 Synergy high-speed cell sorter. The following antibodies were used: anti-mouse CD31 (clone MEC 13.3; BD Biosciences), anti-mouse CD45 (clone 30-F11; BD Biosciences), anti-mouse EpCAM (clone G8.8; BioLegend), anti-AQP3 (Abcam). Anti-AQP3 antibody was labeled using the Zenon antibody labeling kit (Molecular Probes). FACS results were plotted and analyzed using FlowJo software (TreeStar).

### Tissue histology and staining

Lung tissue was fixed with 10% formalin, embedded in paraffin, cut into 5-μm sections and adhered to charged slides. Sections were stained with PAS and hematoxylin as described previously (14, 36). For immunostaining, sections were deparaffinized in Fisherbrand® CitriSolv® (Fisher), hydrated, and heat-treated with antigen unmasking solution (Vector Laboratories, Inc). Immunostaining was performed with the commercially available primary antibodies detailed in **Supplemental Table 3** and a rabbit anti-CLCA1 antibody described previously (1). Primary Abs were detected with secondary Abs labeled with Alexa Fluor 488 (Thermo Fisher Scientific) or Alexa Fluor 594 (Thermo Fisher Scientific) followed by DAPI counterstaining. Slides were imaged by light microscopy using a Leica DM5000 B and by immunofluorescent microscopy using an Olympus BX51, and staining was quantified in whole lung sections using a NanoZoomer S60 slide scanner (Hamamatsu) and ImageJ software as described previously (14, 36).

### RNA analysis

RNA was purified from homogenized lung tissue using Trizol (Invitrogen) or from isolated cells with the RNeasy mini kit (Qiagen) and was used to generate cDNA with the High-Capacity cDNA Archive kit (Life Technologies). We quantified target mRNA and viral RNA levels using real-time qPCR assay with specific fluorogenic probe-primer combinations and Fast Universal PCR Master Mix systems (Applied Biosystems) with mouse-specific forward and reverse primers and probes as described previously (37) and in **Supplemental Table 4**. For the *Mapk13* assay, primers and probe were designed to amplify WT but not *Mapk13* indel sequence in *Mapk13*^−/−^ mice. All samples were assayed using the 7300HT or QuantStudio 6 Fast Real-Time PCR System and analyzed using Fast System Software (Applied Biosystems). All real-time PCR data was normalized to the level of *Gapdh* mRNA. Values were expressed as fold-change based on the delta-delta Ct method as described previously (73).

### Lung function tests

To test lung function, we assessed blood oxygen saturation (SpO_2_) and airway reactivity to methacholine (MCh) as described previously (14). For SpO_2_ monitoring, we used a MouseSTAT® pulse oximeter (Kent Scientific, Torrington, CT) applied to the paw skin under isoflurane anesthesia as described previously (14). To decrease variability in readings for this procedure, anesthetized mice were placed on the RightTemp warming pad, and the SpO_2_ reading was recorded when the Oxiwave signal was stable. For airway reactivity measurements, the response to inhaled methacholine was determined by monitoring respiratory system resistance (R_RS_) as described previously (14).

### Human clinical samples

For asthma, COPD, and non-disease control samples, lung tissue was obtained from our Advanced Lung Disease Tissue Registry that contains whole lung explants harvested from consented patients in our lung transplant program; lungs that were not usable for transplantation from the local Organ Procurement Organization, Mid-America Transplant; and lungs from a tissue procurement service (IIAM, Edison, NJ) as described previously (1, 4, 74, 75). Human studies were conducted with protocols approved by the Washington University (St. Louis, MO) Institutional Review Board and USAMRDC Office of Research Protections. Clinical characteristics of tissue donors are provided in **Supplemental Table 1**.

### Mouse and human airway epithelial cell-derived organoids

For mouse airway organoids, lung epithelial cells from FACS (as described above and previously [Wu 2024)) were resuspended in SAGM (Lonza) mixed 1:1 with growth-factor reduced Matrigel (BD Biosciences) and plated at 3 x 10^3^ cells per 100 μL in 24-well Transwells (Corning). For the first 5 d, SAGM was added with 10 µM Rock inhibitor (Sigma) but without FBS. After 5 d, SAGM was added with 10 % FBS but without Rock inhibitor. The medium was changed every 2-3 d. Organoids were cultured with media alone for 14 d to monitor cell stemness (using organoid numbers and size) and with media plus IL-13 to monitor mucous metaplasia (using *Muc5ac* mRNA) as described previously (42). For human airway organoids, human tracheal and bronchial epithelial cells (hTECs) were isolated by enzymatic digestion and cultured under organoid-forming conditions as described previously [Wu 2024]. For the present experiments, hTECs were cultured in 96-well plate (2 x 10^4^ cells per well) with two distinct AUMsilence FANA oligonucleotides specific for *Mapk13* mRNA and a scrambled control (AUM BioTech), with each ASO added to cells at 1 µM final concentration on culture days 1 and 3. MAPK13 protein level was assessed using western blot as described above. Organoid formation and mucous metaplasia were assessed as described for the corresponding mouse model.

### Statistical analysis

All data are presented as mean and s.e.m. and are representative of at least three experiments with at least 5 data points per experiment. For cell and molecular data, unpaired student’s t-test with Bonferroni correction as well as mixed-model repeated measures analysis of variance with Tukey correction for multiple comparisons were used to assess statistical significance between means. In all cases, significance threshold was set at *P* < 0.05. For physiological data, airway reactivity was assessed using either two-way repeated measures analysis of variance or restricted maximum likelihood linear mixed model with genotype, infection, and methacholine dose as fixed effects and subject as a random effect.

## Abbreviations used in this article

basal-ESC: basal-epithelial stem cell
CLCA1: chloride channel accessory 1
COPD: chronic obstructive pulmonary disease
hTEC: human tracheobronchial epithelial cell
MAPK: mitogen-activated protein kinase
MUC5AC: mucin 5AC
MUC5B: mucin 5B
SeV: Sendai virus.

## Supplemental Materials

Supplemental Figure 1 and Tables 1-4.

## Acknowledgments

We thank the Pulmonary Morphology Core, Pathology-Immunology Transgenic-Microinjection Core, Genome Technology Access Core, Genome Engineering and iPSC Center, Anatomic and Molecular Pathology Core Labs, and Division of Comparative Medicine for technical support.

## Author contributions

K.W. performed mouse, cell, and human tissue experiments; Y.Z. generated *Mapk13*^−/−^ mice and performed mouse and cell experiments; D.M. performed mouse and cell experiments; C.A.I performed mouse and cell experiments; H.Y-D. performed immunostaining experiments; K.S. performed mouse experiments; S.P.K. performed mouse and cell experiments; H.A.W. performed cell experiments; D.Y. performed mouse experiments; J.Y. performed cell experiments, S.R.A. obtained and registered human samples; D.E.B. obtained and registered human samples, S.L.B. generated human epithelial cell samples, .E.C.C. analyzed histology; A.G.R. analyzed kinase-related data;, and M. J. H. directed the project and wrote the manuscript.

## References

1. Alevy Y, Patel AC, Romero AG, Patel DA, Tucker J, Roswit WT, Miller CA, Heier RF, Byers DE, Brett TJ, and Holtzman MJ. IL-13–induced airway mucus production is attenuated by MAPK13 inhibition. J Clin Invest. 2012;122:4555–68.

2. Kim EY, Battaile JT, Patel AC, You Y, Agapov E, Grayson MH, Benoit LA, Byers DE, Alevy Y, Tucker J, Swanson S, Tidwell R, Tyner JW, Morton JD, Castro M, Polineni D, Patterson GA, Schwendener RA, Allard JD, Peltz G, and Holtzman MJ. Persistent activation of an innate immune response translates respiratory viral infection into chronic lung disease. Nat Med. 2008;14:633–40.

3. Byers DE, Alexander-Brett J, Patel AC, Agapov E, Dang-Vu G, Jin X, Wu K, You Y, Alevy YG, Girard J-P, Stappenbeck TS, Patterson GA, Pierce RA, Brody SL, and Holtzman MJ. Long-term IL-33-producing epithelial progenitor cells in chronic obstructive lung disease. J Clin Invest. 2013;123:3967–82.

4. Wu K, Zhang Y, Yin Declue H, Austin SR, Byers DE, Crouch EC, and Holtzman MJ. Lung remodeling regions in long-term coronavirus disease 2019 feature basal epithelial cell reprogramming. Am J Pathol 2023;193:680–9.

5. Dunican EM, Elicker BM, Gierada DS, Nagle SK, Schiebler ML, Newell JD, Raymond WW, Lachowicz-Scroggins ME, Di Maio S, Hoffman EA, Castro M, Fain SB, Jarjour NN, Israel E, Levy BD, Erzurum SC, Wenzel SE, Meyers DA, Bleecker ER, Phillips BR, Mauger DT, Gordon ED, Woodruff PG, Peters MC, and Fahy JV. Mucus plugs in patients with asthma linked to eosinophilia and airflow obstruction. J Clin Invest. 2018;128:997–1009.

6. Toda M, Tulic MK, Levitt RC, and Hamid Q. A calcium-activated chloride channel (HCLCA1) is strongly related to IL-9 expression and mucus production in bronchial epithelium of patients with asthma. J Allergy Clin Immunol. 2002;109(2):246–50.

7. Tang M, Elicker BM, Henry T, Gierada DS, Schiebler ML, Huang BK, Peters MC, Castro M, Hoffman EA, Fain SB, Ash SY, Choi JY, Hall C, Phillips BR, Mauger DT, Denlinger LC, Jarjour NN, Israel E, Phipatanakul W, Levy BD, Wenzel SE, Bleecker ER, Woodruff PG, Fahy JV, Dunican EM, and Program-3 NSAR. Mucus plugs persist in asthma, and changes in mucus plugs associate with changes in airflow over time. Am J Respir Crit Care Med. 2022;205:1036–45.

8. Hogg JC, Chu F, Utokaparch S, Woods R, Elliott WM, Buzatu L, Cherniack RM, Rogers RM, Sciurba FC, Coxson HO, and Pare PD. The nature of small-airway obstruction in chronic obstructive pulmonary disease. N Eng J Med. 2004;350:2645–53.

9. Ekberg-Aronsson M, Pehrsson K, Nilsson JA, Nilsson PM, and Lofdahl CG. Mortality in GOLD stages of COPD and its dependence on symptoms of chronic bronchitis. Respir Res. 2005;6:98.

10. Hogg JC, Chu FS, Tan WC, Sin DD, Patel SA, Pare PD, Martinez FJ, Rogers RM, Make BJ, Criner GJ, Cherniack RM, Sharafkhaneh A, Luketich JD, Coxson HO, Elliott WM, and Sciurba FC. Survival after lung volume reduction in chronic obstructive pulmonary disease: insights from small airway pathology. Am J Respir Crit Care Med. 2007;176(5):454–9.

11. Kesimer M, Ford AA, Ceppe A, Radicioni G, Cao R, Davis CW, Doerschuk CM, Alexis NE, Anderson WH, Henderson AG, Barr RG, Bleecker ER, Christenson SA, Cooper CB, Han MK, Hansel NN, Hastie AT, Hoffman EA, Kanner RE, Martinez F, Paine Rr, Woodruff PG, O’Neal WK, and Boucher RC. Airway mucin concentration as a marker of chronic bronchitis. N Engl J Med. 2017;377:911–22.

12. Zuo W, Zhang T, Zheng D, Wu A, Guan SP, Liew A-A, Yamamoto Y, Wang X, Lim SJ, Vincent M, Lessard M, P. CC, Xian W, and McKeon F. p63+Krt5+ distal airway stem cells are essential for lung regeneration. Nature. 2015;517:616–20.

13. Vaughan AE, Brumwell AN, Xi Y, Gotts JE, Brownfield DG, Treutlein B, Tan K, Tan V, Liu FC, Looney MR, Matthay MA, Rock JR, and Chapman HA. Lineage-negative progenitors mobilize to regenerate lung epithelium after major injury. Nature. 2015;517:621–5.

14. Wu K, Kamimoto K, Zhang Y, Yang K, Keeler SP, Gerovac BJ, Agapov EV, Austin SP, Yantis J, Gissy KA, Byers DE, Alexander-Brett J, Hoffmann CM, Wallace M, Hughes ME, Morris SA, and Holtzman MJ. Basal-epithelial stem cells cross an alarmin checkpoint for post-viral lung disease. J Clin Invest. 2021;131:e149336.

15. Taylor MS, Chivukula RR, Myers LC, Jeck WR, Waghray A, Tata PR, Selig MK, O’Donnell WJ, Farver CF, Thompson BT, Rajagopal J, and Kradin RL. A conserved distal lung regenerative pathway in acute lung injury. Am J Pathol. 2018;188:1149–60.

16. Delorey TM, Ziegler CGK, Heimberg G, Normand R, Yang Y, Segerstolpe A, Abbonddanza D, Fleming SJ, Subramanian A, Montoro DT, Jagadeesh KA, Dey KD, Sen P, Slyper M, Pita-Juarez YH, and Phillips D. COVID-19 tissue atlases reveal SARS-CoV-2 pathology and cellular targets. Nature. 2021;595:107–13.

17. Holtzman MJ. Asthma as a chronic disease of the innate and adaptive immune systems responding to viruses and allergens. J Clin Invest. 2012;122:2741–8.

18. Han J, Wu J, and Silke J. An overview of mammalian p38 mitogen-activated protein kinases, central regulators of cell stress and receptor signaling. F1000Res. 2020;9.

19. Haller V, Nahidino P, Forster M, and Laufer SA. An updated patent review of p38 MAPK kinase inhibitors (2014-2019). Expert Opin Ther Pat. 2020;30:453–66.

20. Pelaia C, Vatre, A., Gallelli L, Lombardo N, Sciacqua A, Savino R, and Pelaia G. Role of p38 mitogen-activated protein kinase in asthma and COPD: pathogenic aspects and potential targeted therapies. Drug Design, Development and Therapy. 2021;15:1275–84.

21. Martucci C, Allen AD, Moretto N, Bagnacani V, Fioni A, Patacchini R, Civelli M, Villetti G, and Facchinetti F. CHF6297: a novel potent and selective p38 MAPK inhibitor with robust anti-inflammatory activity and suitable for inhaled pulmonary administration as dry powder. Front Pharmacol. 2024;15:1343941.

22. Aikawa T, Shimura S, Sasaki H, Ebina M, and Takishima T. Marked goblet cell hyperplasia with mucus accumulation in the airways of patients who died of severe acute asthma attack. Chest. 1992;101:916–21.

23. Radicioni G, Ceppe A, Ford AA, Alexis NE, Barr RG, Bleecker ER, Christenson SA, Cooper CB, Han MK, Hansel NN, Hastie AT, Hoffman EA, Kanner RE, Martinez FJ, Ozkan E, Paine R, 3rd, Woodruff PG, O’Neal WK, Boucher RC, and Kesimer M. Airway mucin MUC5AC and MUC5B concentrations and the initiation and progression of chronic obstructive pulmonary disease: an analysis of the SPIROMICS cohort. Lancet Respir Med. 2021;9(11):1241–54.

24. Goedert M, Hasegawa M, Jakes R, Lawler S, Cuenda A, and Cohen P. Phosphorylation of microtube-associated protein tau by stress-activated protein kinases. FEBS Lett. 1997;409:57–62.

25. Sumara G, Formentini I, Collins S, Sumara I, Windak R, Bodenmiller B, Ramracheya R, Caille D, Jiang H, Platt KA, Meda P, Aebersold R, Rorsman P, and Ricci R. Regulation of PKD by the MAPK p38δ in insulin secretion and glucose homeostasis. Cell. 2009;136:235–48.

26. Ittner A, Block H, Reichel CA, Varjosalo M, Gehart H, Sumara G, Gstaiger M, Krombach F, Zarbock A, and Ricci R. Regulation of PTEN activity by p38d-PKD1 signaling in neutrophils confers inflammatory responses in the lung. J Exp Med. 2012;209:2229–46.

27. Schindler E, Hindes A, Gribben E, Burns C, Yin Y, Lin M, Owen R, Longmore G, Kissling G, Arthur J, and Efimova T. p38delta Mitogen-activated protein kinase is essential for skin tumor development in mice. Cancer Res. 2009;69:4648–55.

28. Risco A, Del Fresno C, Mambol A, Alsina-Beauchamp D, MacKenzie KF, Yang H-T, Barber DF, Morcelle C, Arthur JSC, Ley SC, Ardavin C, and Cuenda A. p38g and p38d kinases regulate the Toll-like receptor 4 (TLR4)-induced cytokine production by controlling ERK1/2 protein kinase pathway activation. Proc Natl Acad Sci U S A. 2012;109:11200–5.

29. Tomas-Loba A, Manieri E, Gonzalez-Teran B, Mora A, Leiva-Vega L, Santamans AM, Romero-Becerra R, Rodriguez E, Pintor-Chocano A, Feixas F, Lopez JA, Caballero B, Trakala M, Blanco O, Torres JL, Hernandez-Cosido L, Montalvo-Romeral V, Matesanz N, Roche-Molina M, A. BJ, Mischo H, Leon M, Caballero A, Miranda-Saavedra D, Ruiz-Cabello J, Nevzorova YA, Cubero FJ, Bravo J, Vazquez J, Malumbres M, Marcos M, Osuna S, and Sabio G. p38g is essential for cell cycle progression and liver tumoriogenesis. Nature. 2019;568:557–60.

30. Tyner JW, Kim EY, Ide K, Pelletier MR, Roswit WT, Morton JD, Battaile JT, Patel AC, Patterson GA, Castro M, Spoor MS, You Y, Brody SL, and Holtzman MJ. Blocking airway mucous cell metaplasia by inhibiting EGFR antiapoptosis and IL-13 transdifferentiation signals. J Clin Invest. 2006;116:309–21.

31. Patel AC, Morton JD, Kim EY, Alevy Y, Swanson S, Tucker J, Huang G, Agapov E, Phillips TE, Fuentes ME, Iglesias A, Aud D, Allard JD, Dabbagh K, Peltz G, and Holtzman MJ. Genetic segregation of airway disease traits despite redundancy of chloride channel calcium-activated (CLCA) family members. Physiol Genomics. 2006;25:502–13.

32. Grayson MH, Cheung D, Rohlfing MM, Kitchens R, Spiegel DE, Tucker J, Battaile JT, Alevy Y, Yan L, Agapov E, Kim EY, and Holtzman MJ. Induction of high-affinity IgE receptor on lung dendritic cells during viral infection leads to mucous cell metaplasia. J Exp Med. 2007;204:2759–69.

33. Agapov E, Battaile JT, Tidwell R, Hachem R, Patterson GA, Pierce RA, Atkinson JJ, and Holtzman MJ. Macrophage chitinase 1 stratifies chronic obstructive lung disease. Am J Respir Cell Mol Biol. 2009;41:379–84.

34. Wu K, Byers DE, Jin X, Agapov E, Alexander-Brett J, Patel AC, Cella M, Gilfilan S, Colonna M, Kober DL, Brett TJ, and Holtzman MJ. TREM-2 promotes macrophage survival and lung disease after respiratory viral infection. J Exp Med. 2015;212:681–97.

35. Keeler SP, Agapov EV, Hinojosa ME, Letvin AN, Wu K, and Holtzman MJ. Influenza A virus infection causes chronic lung disease linked to sites of active viral RNA remnants. J Immunol. 2018;201:2354–68.

36. Zhang Y, Mao D, Keeler SP, Wang X, Wu K, Gerovac BJ, Shornick LP, Agapov E, and Holtzman MJ. Respiratory enterovirus (like parainfluenza virus) can cause chronic lung disease if protection by airway epithelial STAT1 is lost. J Immunol. 2019;202:2332–47.

37. Wu K, Wang X, Keeler SP, Gerovac BJ, Agapov E, Byers DE, Gilfillan S, Colonna M, Zhang Y, and Holtzman MJ. Group 2 innate lymphoid cells must partner with the myeloid-macrophage lineage for long-term postviral lung disease. J Immunol. 2020;205:1084–101.

38. Wang X, Wu K, Keeler SP, Mao D, Agapov EV, Zhang Y, and Holtzman MJ. TLR3-activated monocyte-derived dendritic cells trigger progression from acute viral infection to chronic disease in the lung. J Immunol. 2021;206:1297–314 (selected for Top-Reads p. 115).

39. Keeler SP, Yantis J, Youkilis S, Podgorny S, Mao D, Zhang Y, Whitworth KM, Redel BK, Samuel MS, Wells KD, Prather RS, and Holtzman MJ. Chloride channel accessory 1 (CLCA1) gene deficiency causes selective loss of mucus production in a new pig model. Am J Physiol Lung Cell Mol Physiol. 2022;322:L842–L52.

40. Martin RA, Keeler SP, Wu K, Shearon WJ, Patel D, Hoang M, Hoffmann CM, Hughes ME, and Holtzman MJ. An alternative mechanism for skeletal muscle dysfunction in long-term post-viral lung disease. bioRxiv. 2022;10.1101/2022.10.07.511313.

41. Walter MJ, Morton JD, Kajiwara N, Agapov E, and Holtzman MJ. Viral induction of a chronic asthma phenotype and genetic segregation from the acute response. J Clin Invest. 2002;110:165–75.

42. Wu K, Zhang Y, Yin-Declue H, Sun K, Mao D, Austin S, Crouch E, Brody S, Byers D, Hoffmann C, Hughes M, and Holtzman M. A correctable immune niche for basal-epithelial stem cell reprogramming and post-viral lung diseases. J Clin Invest. 2024;134:e183092.

43. Wu K, Zhang Y, Yin Declue H, Sun K, Mao D, Crouch EC, Byers DE, and Holtzman MJ. The post-viral GPNMB^+^ immune niche persists in long-term Covid, asthma, and COPD. medRxiv. 2024;2024.08.27.24312640:doi: 10.1101/2024.08.27.24312640.

44. Keeler SP, Wu K, Zhang Y, Mao D, Li M, Iberg CA, Austin SP, Glaser SA, Yantis J, Podgorny S, Brody SL, Chartock JR, Han Z, Byers DE, Romero AG, and Holtzman MJ. A potent MAPK13-14 inhibitor prevents airway inflammation and mucus production. Am J Physiol Lung Cell Mol Physiol. 2023;325:L726–40.

45. Holtzman MJ, Zhang Y, Wu K, and Romero AG. Mitogen-activated protein kinase-guided drug discovery for post-viral and related types of lung disease. Eur Respir Rev. 2024;33:230220.

46. Sabio G, Simon J, Arthur C, Kuma Y, Peggie M, Carr J, Murray-Tait V, Centeno F, Goedert M, Morrice NA, and Cuenda A. p38g regulates the localisation of SAP97 in the cytoskeleton by modulating its interaction with GKAP. EMBO J. 2005;24:1134–45.

47. Zur R, Garcia-Ibanez L, Nunez-Buiza A, Aparicio N, Liappas G, Escos A, Risco A, Page A, Saiz-Ladera C, Alsina-Beauchamp D, Montans J, Paramio JM, and Cuenda A. Combined deletion of p38gamma and p38delta reduces skin inflammation and protects from carcinogenesis. Oncotarget. 2015;6(15):12920–35.

48. Gonzalez-Teran B, Lopez JA, Rodriguez E, Leiva L, Martinez-Martinez S, Bernal JA, Jimenez-Borreguero LJ, Redondo JM, Vazquez J, and Sabio G. p38gamma and delta promote heart hypertrophy by targeting the mTOR-inhibitory protein DEPTOR for degradation. Nat Commun. 2016;7:10477.

49. Alsina-Beauchamp D, Escos A, Fajardo P, Gonzalez-Romero D, Diaz-Mora E, A. R, Martin-Serrano MA, C. dF, Dominguez-Andres J, Aparicio N, Zur R, Shpiro N, Brown GD, Ardavin C, Netea MC, Alemany S, Sanz-Ezquerro JJ, and Cuenda A. Myeloid cell deficiency of p38g/p38d protects against candidiasis and regulates antifungal immunity. EMBO Mol Med. 2018;10:e8485.

50. Barrio L, Roman-Garcia S, Diaz-Mora E, Risco A, Jimenez-Saiz R, Carrasco YR, and Cuenda A. B Cell Development and T-Dependent Antibody Response Are Regulated by p38gamma and p38delta. Front Cell Dev Biol. 2020;8:189.

51. Escos A, Martin-Gomez J, Gonzalez-Romero D, Diaz-Mora E, Francisco-Velilla R, Santiago C, Cueza J, Dominguez-Zorita S, Martinez-Salas E, Sonenberg N, Sanz-Ezquerro J, Jafanejad S, and Cuenda A. TPL2 kinase expression is regulated by the p38g/p38d-dependent association with aconitase-1 with *TPL2* mRNA. Proc Natl Acad Sci U S A. 2022;119:e2204752119.

52. Escos A, Diaz-Mora E, Pattison M, Fajardo P, Gonzalez-Romero D, Risco A, Martin-Gomez J, Bonnell E, Sonenberg N, Jafanejad SM, Sanz-Ezquerro JJ, Ley SC, and Cuenda A. p38g and p38d modulate innate immune response by regulating MEF2D activation. eLife. 2023;12:e86200.

53. Underwood DC, Osborn RR, and Kotzer CJ. SB 239063, a potent p38 MAP kinase inhibitor, reduces inflammatory cytokine production, airways eosinophil infiltration, and persistence. J Pharmacol Exp Ther. 2000;293:281–8.

54. Duan W, Chan JH, McKay K, Crosby JR, Choo HH, Leung BP, Karras JG, and Wong WSF. Inhaled p38alpha mitogen-activated protein kinase antisense oligonucleotide attenuates asthma in mice. Am J Respir Crit Care Med. 2005;171:571–8.

55. Medicherla S, Fitzgerald MF, Spicer D, Woodman P, Ma JY, Kapoun AM, Chakravarthy S, Dugar S, Protter AA, and Higgins LS. p38a-selective mitogen-activated protein kinase inhibitor SD-282 reduces inflammation in a subchronic model of tobacco smoke-induced airway inflammation. J Pharmacol Exp Ther. 2008;324:921–9.

56. Renda T, Baraldo S, Pelaia G, Bazzan E, Turato G, Papi A, Maestrelli P, Maselli R, Vatrella A, Fabbri LM, Zuin R, Marsico SA, and Saetta M. Increased activation of p38 MAPK in COPD. Eur Respir J. 2008;31:62–9.

57. MacNee W, Allan RJ, Jones I, Cristina De Salvo M, and Tan LF. Efficacy and safety of the oral p38 inhibitor PH-797804 in chronic obstructive pulmonary disease: a randomised clinical trial. Thorax. 2013;68:738–45.

58. Kawai T, and Akira S. Pathogen recognition with Toll-like receptors. Curr Opin Immunol. 2005;17(4):338–44.

59. Ha U, H. LJ, Jono H, Koga T, Srivastava A, Malley R, Pages G, Pouyssegur J, and Li J-D. A novel role for IkB kinase (IKK) a and IKKb in ERK-dependent up-regulation of MUC5AC mucin transcription by Streptococcus pneumoniae. J Immunol. 2007;178:1736–47.

60. Fujisawa T, Velichko S, Thai P, Hung LY, Huang F, and Wu R. Regulation of airway MUC5AC expression by IL-1beta and IL-17A; the NF-kappaB paradigm. J Immunol. 2009;183(10):6236–43.

61. Na HG, Kim Y-D, and Bae CH. High concentration of insulin induces MUC5AC expression via phosphoinositide 3 kinase/AKT and mitogen-activated protein kinase signaling pathways in human airway epithelial cells. Am J Rhinol Allergy. 2018.

62. Xu H, Sun Q, Lu L, Luo F, Zhou L, Liu J, Cao L, Wang Q, Xue J, Yang Q, Yang P, Lu J, Xiang Q, and Liu Q. MicroRNA-218 acts by repressing TNFR1-mediated activation of NF-kB, which is involved in MUC5AC hyper-production and inflammation in smoking-induced bronchiolitis of COPD. Toxicol Lett. 2017;280:171–80.

63. Wu S, Li H, Yu L, Wang N, Li X, and Chen W. IL-1b upregulates Muc5ac expression via NF-kB-induced HIF-1a in asthma. Immunol Lett. 2017;192:20–6.

64. Selness SR, Devraj RV, Devadas B, Walker JK, Boehm TL, Durley RC, Shieh H, Xing L, Rucker PV, Jerome KD, and Benson AG. Discovery of PH-797804, a highly selective and potent inhibitor of p38 MAP kinase. Bioorg Med Chem Lett. 2011;21:4066–71.

65. McQualter JL, Yuen K, Williams B, and Bertoncello I. Evidence of an epithelial stem/progenitor cell hierarchy in the adult mouse lung. Proc Natl Acad Sci U S A. 2010;107(4):1414–9.

66. Tarutani M, Itami S, Okabe M, Ikawa M, Tezuka T, Yoshikawa K, Kinoshita T, and Takeda J. Tissue-specific knockout of the mouse Pig-a gene reveals important roles for GPI-anchored proteins in skin development. American Review of Respiratory Disease. 1997;94:7400–5.

67. Varjosalo M, Keskitalo S, Van Drogen A, Nurkkala H, Vichalkovski A, Aebersold R, and Gstaiger M. The protein interaction landscape of the human CMGC kinase group. Cell Rep. 2013;3(1306-1320).

68. Zhang Y, Wu K, Mao D, Iberg CA, Yin-Declue H, Sun K, Wikfors H, Keeler SP, Li M, Young D, Yantis J, Austin S, Crouch E, Chartock J, Han Z, Byers D, Brody S, Romero AG, and Holtzman M. A first-in-kind MAPK13 inhibitor corrects stem cell reprogramming and muco-obstructive lung disease. bioRxiv. 2025;10.1101/2024.08.21.608990.

69. Sigurs N, Aljassim F, Kjellman B, Robinson PD, Sigurbergsson F, Bjarnason R, and Gustafsson PM. Asthma and allergy patterns over 18 years after severe RSV bronchiolitis in the first year of life. Thorax. 2010;65(12):1045–52.

70. Holtzman MJ, Byers DE, Alexander-Brett J, and Wang X. The role of airway epithelial cells and innate immune cells in chronic respiratory disease. Nat Rev Immunol. 2014;14:686–98.

71. Mallia P, Message SD, Gielen V, Contoli M, Gray K, Kebadze T, Aniscenko J, Laza-Stanca V, Edwards MR, Slater L, Papi A, Stanciu LA, Kon OM, Johnson M, and Johnston SL. Experimental rhinovirus infection as a human model of chronic obstructive pulmonary disease exacerbation. Am J Respir Crit Care Med. 2011;183:734–42.

72. van Nunen MCJ, and van der Veen J. Experimental infection with Sendai virus in mice. Arch Gesamte Virusforsch. 1967;22:388–97.

73. Livak KJ, and Schmittgen TD. Analysis of relative gene expression data using real-time quantitative PCR and the 2(-Delta Delta C(T)) Method. Methods. 2001;25:402–8.

74. Deslee G, Woods J, Moore C, Conradi S, Gierada D, Atkinson J, Battaile J, Liu L, Patterson A, Adair-Kirk T, Holtzman M, and Pierce R. Oxidative damage to nucleic acids in severe emphysema. Chest. 2009;135:965–74.

75. Byers DE, Wu K, Dang-Vu G, Jin X, Agapov E, Zhang X, Battaile JT, Schechtman KB, Yusen R, Pierce RA, and Holtzman MJ. Triggering receptor expressed on myeloid cells-2 (TREM-2) expression tracks with M2-like macrophage activity and disease severity in COPD. Chest. 2018;153:77–86.

