## Supplemental Materials for "MAPK13 controls structural remodeling and disease after epithelial injury"

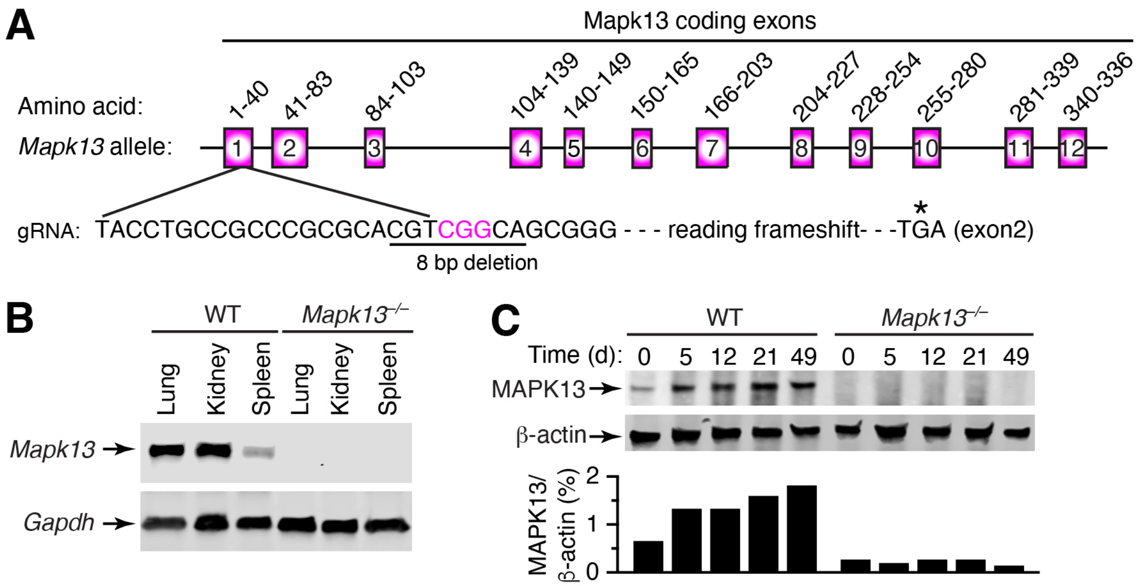


**Supplemental Fig. 1. Design and validation of *Mapk13* gene knockout (*Mapk13*$^{-/-}$) mice. A,** Design for generation of *Mapk13*$^{-/-}$ mice using genetically susceptible C57BL/6J strain. **B,** Western blot for levels of MAPK13 and β-actin control for indicated tissues from WT and *Mapk13*$^{-/-}$ mice under baseline conditions. **C,** Western blot for levels of MAPK13 and β-actin control for lung tissue from WT and *Mapk13*$^{-/-}$ mice at 0-49 d after SeV infection along with corresponding quantitation.

**Supplemental Table 1. Characteristics of tissue donor groups.**

| Characteristic | Donor (n=11) | Asthma (n=10) | COPD (n=10) |
|---|---|---|---|
| Age | 43 ± 18.4 | 43.8 ± 17.6 | 61.8 ± 5.7 |
| Sex (M/F) | 7/4 | 5/5 | 4/6 |
| Race (W/B/H) | 7/0/0[1] | 4/4/2 | 9/1/0 |
| Pack-year | ND[2] | None[3] | 48.5 ± 27.0 |
| Years quit | ND | ND | 10.3 ± 8.8 |
| FVC % pred | ND | ND | 67.5 ± 11.4 |
| FEV1 % pred | ND | ND | 21.0 ± 4.1 |
| FEV1/FVC ratio | ND | ND | 0.25 ± 0.04 |

[1]Race data missing for some donor subjects.

[2]Two deceased donor subjects had a history of tobacco smoking 10 pack-years and 15 pack-years, but smoking data was not available for all subjects.

[3]Two deceased asthmatics had less than a 5 pack-yr history of tobacco smoking.

Abbreviations: ND, not determined.

**Supplemental Table 2. Sequences of DNA primers and probes for determining levels of SeV RNA in real-time qPCR assays.**

| Target gene | Type | Sequence |
|---|---|---|
| SeV-*NP* | F[1] | 5'-GGCGGTGGTGCAATTGAG-3' |
| | R | 5'-CATGAGCTTCTGTTTCTAGGTCGAT-3' |
| | P | 5'-AGCTCTAGACAATGCC-3' |

[1]Abbreviations: F, forward primer; R, reverse primer; P, MGB probe.

**Supplemental Table 3. Antibodies for immunostaining mouse and human tissues.**

| Target Protein | Antibody Type | Vendor | Catalogue # |
|---|---|---|---|
| F4/80 | Rabbit mAb | Cell Signaling | 70076 |
| Mouse Il33 | Goat pAb | R&D Systems | AF3626 |
| Human Il33 | Rabbit pAb | Sigma | HPA024426 |
| Ki-67 | Mouse mAb | BD Pharmingen | 550609 |
| Ki-67 | Rabbit mAb | Cell Signaling | 12202 |
| Krt5 | Rabbit pAb | Abcam | ab53121 |
| Krt5 | Chicken pAb | Biolegend | 905904 |
| Mapk13/MAPK13 | Rabbit pAb | R&D Systems | AF1519 |
| MBP | Mouse mAb | Fisher | NBP1421401M |
| Muc5ac | Mouse mAb (45M1), biotinylated | Thermo Scientific | MS-145-B |
| Muc5ac | Mouse mAb (45M1) | Thermo Scientifc | MS-145-P |
| Muc5b | Rabbit pAb | Abcam | ab87276 |
| Nos2 | Rabbit pAb | Abcam | ab3523 |
| Sftpc | Rabbit pAb | Abcam | ab90716 |

[1]Abbreviations: mAb, monoclonal antibody, pAb, polyclonal antibody.

**Supplemental Table 4. Sequences of DNA primers and probes for qPCR assays in mouse tissue.**

| Target Gene | Type | ID/Sequence |
|---|---|---|
| *Arg1* | | Mm00475988_m1 (ThermoFisher Scientific) |
| *Aqp3* | | Mm.PT.58.13308206 (Integrated DNA Technologies) |
| *Clca1* | F[1]<br>R<br>P | 5'-ACCGGCTGCCGCTAAAGAGCTTGAG-3'<br>5'-AGACCATTGTTCTGAACCTGATCCGAAG-3'<br>5'-AGCTGTCCAAAATGACAGGAGGCCTGCAGACATA-3' |
| *Cxcl17* | | Mm.PT.58.28640067 (Integrated DNA Technologies) |
| *Gapdh* | | Mm.PT.39a.1 (Integrated DNA Technologies) |
| *IFNg* | | Mm.PT.58.41769240 (Integrated DNA Technologies) |
| *Il1b* | | Mm.PT.58.41616450 (Integrated DNA Technologies) |
| *Il6* | | Mm.PT.58.10005566 (Integrated DNA Technologies) |
| *Il13* | F<br>R<br>P | 5'-GGAGCTGAGCAACATCACACA-3'<br>5'-CACACTCCATACCATGCTGCC-3'<br>5'-CCAGACTCCCCTGTGCA-3' |
| *Il33* | | Mm00505403_m1 (ThermoFisher Scientific) |
| *Krt5* | | Mm.PT.58.41573083 (Integrated DNA Technologies) |
| *Ltf* | | Mm00434787_m1 (ThermoFisher Scientific) |
| *Mapk13* | F<br>R<br>P | 5'-GGAGCTACCCAAGACCTACCT-3'<br>5'-TGTCCGCTTGTCGATGGCCGA-3'<br>5'-GCGCACGTCGGCA-3' |
| *Muc5ac* | F<br>R<br>P | 5'-TACCACTCCCTGCTTCTGCAGCGTGTCA-3'<br>5'-ATAGTAACAGTGGCCATCAAGGTCTGTCT-3'<br>5'-TATACCCCTTGGGATCCATCATCTACA-3' |
| *Muc5b* | F<br>R<br>P | 5'-CTTTCACCCTCAGGAACACGAT-3'<br>5'-TTCGAGGATTATACAGTTCAAAGCA-3'<br>5'-TGAAGGACAAGGTGTGGAGATT-3' |
| *Nos2* | | Mm.PT.58.43705194 (Integrated DNA Technologies) |
| *SerpinB2* | | Mm.PT.58.13584177 (Integrated DNA Technologies) |
| *Tnfa* | | Mm.PT.58.12575861 (Integrated DNA Technologies) |
| *Trem2* | | Mm.PT.58.7992121 (Integrated DNA Technologies) |
| *Trp63* | | Mm.PT.58.11081628 (Integrated DNA Technologies) |

[1]Abbreviations: F, forward primer; R, reverse primer; P, MGB probe.
